# Alzheimer disease and Apolipoprotein E4: meningeal brain lymphatics point to new clues in pathogenesis

**DOI:** 10.1101/571729

**Authors:** Alexios-Fotios A. Mentis, George P. Chrousos

## Abstract

The role of the lymphatic system in brain function and/or dysfunction has long been an enigma. However, recent reports that meningeal lymphatic vessels exist within the mouse and human brain, as well as evidence that mouse meningeal lymphatic vessels play a role in clearing the toxic amyloid-beta peptide connected with Alzheimer’s disease (AD), may herald novel diagnostic and therapeutic avenues. Here, we explore new evidence connecting the lymphatic system of the brain with AD. In particular, we focus on new findings showing that meningeal lymphatic vessels play a role in drainage of cerebrospinal fluid and egress of immune cells from the brain, and that disrupting this vessel system leads to accumulation of amyloid - beta peptide and cognitive dysfunction. We also discuss the hypothesis that apolipoprotein E isoform e4 (APO E4) ─ the leading genetic risk for developing AD ─ is involved in meningeal lymphatic vessel function. By reanalyzing previously published RNA-Seq data, we show that APO E4 knock-in microglia cells express lower levels of genes representing lymphatic markers (a phenomenon we call “attenuated lymphaticness”) and of genes in which functional missense mutations are linked to lymphedema. Accordingly, we propose the hypothesis that APO E4 is involved in the shrinkage of lymphatic vessels. This notion could lead, if verified by additional anatomic and mechanistic data, to the concept that APO E4-related AD (such as in late onset AD or trisomy 21-related AD) is related to lymphosclerosis coupled with lymphedema.

## 1. Lymphatic vessels in the brain

In peripheral tissues, the lymphatic system plays a crucial role in the clearance of substances Thus, impaired clearance of various substances has been often linked to abnormalities in lymphatic vessels, such as enlarged vessels or substance leakage. Lymphatic vessels do not always originate from the veins, as shown recently by lineage tracing studies; in development, their progenitors include lymphangioblasts, blood cell progenitors, and hemogenic endothelial cells (Klotz *et al*., 2015; Martinez-Corral *et al*., 2015; Pichol-Thievend *et al*., 2018).

Until recently, it was believed that the brain lacks lymphatic vessels. The Italian scientist Paolo Mascagni first described the brain lymphatic system in the 1800’s, but his conclusions were disputed up until recently (for works spanning all this period, see: (Lukic *et al*., 2003) (Foldi *et al*., 1966) (Prineas, 1979) (Andres *et al*., 1987); (Gausas *et al*., 2007); (Furukawa *et al*., 2008)). In addition, despite the description of Mendelian disorders connecting lymphedema and cognitive impairment (e.g., in Hennekan syndrome (Alders *et al*., 2014)) or non-specific neurological signs (e.g., GLANS syndrome) (Berton *et al*., 2015), no suspicion for the potential presence of brain lymphatics had been raised. However, recently, a number of convincing studies have shown that lymphatic vessels exist within the meninges of the central nervous system (CNS) ((Antila *et al*., 2017; Aspelund *et al*., 2015; Louveau *et al*., 2015) and reviewed in (Sun *et al*., 2018)) ─ a notion suggesting *inter alia* that besides meta-research on reproducibility of current findings, the “mining” of “hidden pearls” can be meaningfully crucial to advance biomedical research.

Absinta et al. coupled MRI techniques with post-mortem pathology analysis to show that meningeal lymphatic vessels exist alongside blood vessels in humans and non-human primates (Absinta *et al*., 2017). More recently, Da Mesquita and colleagues demonstrated a functional role for meningeal lymphatic vessels in mice, that they drain macromolecules from the cerebrospinal (CSF) and interstitial fluids of the CNS into cervical lymph nodes. (Da Mesquita *et al*., 2018b) (Da Mesquita *et al*., 2018a). In particular, ablation of the vessels using a photodynamic drug, Visudyne (verteporfin), resulted in an increase in paravascular CSF molecules, efflux of interstitial fluid (ISF) macromolecules, and impairments in cognitive performance, indicating a functional role in the clearance of macromolecules from the mouse brain. These findings are in accordance with previous studies showing that the CSF’ outflow is conducted primarily through the lymphatic vessels, a process compromised during aging ((Ma *et al*., 2017)).

These findings suggest that dysfunctional clearance of macromolecules from the brain by the meningeal lymphatic vessels ─ and the recently deciphered molecular anatomical connections between meninges and skull (Cai *et al*., 2019) ─ will add to the complexity of meningeal architecture and its role in the decline of cognitive function associated with age and neurodegenerative diseases such as Alzheimer’s disease (AD) (De Mesquita, Louveau, et al., 2018).

## 2. Risk factors for AD

Although the majority of AD cases (98%) are sporadic (Louveau *et al*., 2016), with several genetic risk factors identified, environmental factors including chronic stress may also play a role in AD pathogenesis **(Table 1)** (for example, reviewed in (Bellou *et al*., 2017; Louveau *et al*., 2015). In this context, we have shown that stress could contribute to AD through chronic cortisol actions in brain cell metabolism and increases in inflammatory cytokines, mainly catecholamine-induced interleukin-6, as well as TNF-alpha, interleukin 1, and other cytokines secreted by the accumulated visceral fat (reviewed in (Gassen *et al*., 2017; Nicolaides *et al*., 2015)). Stress also appears to act through increased levels of corticotropin-releasing hormone (CRH), which leads to phosphorylation of tau and its accumulation in a globular form in dendritic and axonal neuronal processes (Le *et al*., 2016b).

**Table 1.** Genetic, environmental, and lifestyle risk factors for Alzheimer’s disease.

| Non-genetic risk factors | Genetic – Chromosomal factors |
| --- | --- |
| Chronic stress (psychological and biological) and inflammation<sup>g</sup> |  |
| Depression at any age and late-life depression<sup>a</sup> | Apolipoprotein-E isoform ε4 and other gene loci<sup>c</sup> |
| Type 2 diabetes mellitus<sup>a</sup> | Amyloid precursor protein (inherited AD form)<sup>d</sup> |
| Frequency of social contacts - loneliness<sup>a</sup> | Presenilin-1 gene (inherited AD form)<sup>d</sup> |
| Benzodiazepines use<sup>a</sup> | Presenilin-2 gene (inherited AD form)<sup>d</sup> |
| Low adherence to Mediterranean diet<sup>b</sup> | Trisomy 21 (Down syndrome)<sup>e</sup> |
| Aging<sup>g</sup> | Immune (epigenetically regulated) response<sup>f</sup> |
| Anemia / Lower hemoglobin levels<sup>h</sup> | Loci in IQCK, ACE, ADAM10, ADAMTS1, and WWOX, and rare variants<sup>i,j</sup> |

**Sources:** **a.** (Bellou *et al.*, 2017); **b.** (Psaltopoulou *et al.*, 2013); **c.** (Verheijen and Sleegers, 2018); **d.** (Lanoiselee *et al.*, 2017); **e.** (Mentis, 2016); **f.** (Gjoneska *et al.*, 2015); **g.** (Gassen *et al.*, 2017); **h.** (Winchester *et al.*, 2018); **i.** (Kunkle *et al.*, 2019); **j.** (Jansen *et al.*, 2019)

Interesting connections between chronic stress and the lymphatic system have been observed. For instance, in mouse cancer models, mice subjected to chronic stress by confinement in small spaces showed changes in the structure of their lymph vasculature and increases in occurrence of metastases through the dissemination of tumors through the lymphatic system (Le *et al*., 2016a). Patients taking beta-blockers, which act by reducing adrenergic effects, had significantly fewer lymph node and distant metastases (Le *et al*., 2016a), most likely because of reduction in adrenaline-mediated signaling events. These findings may harbor potentially broader implications, considering the well-established, far-reaching ties of adrenergic stress with neuro-inflammation (Chrousos, 1995).

## 3. The role of Apo E4 in AD

The human apolipoprotein E is encoded by three isoforms: APO E2, APO E3, and APO E4. Carrying the Apo E4 allele is the chief genetic risk factor for developing AD. While the isoform is present in approximately 13-15% of the population, it is carried by more than 50% of individuals with late-onset AD (Xian *et al*., 2018). An individual possessing a single allele has a threefold increased risk of developing AD late in life, while carrying two alleles has been associated with a higher than 10-fold risk (Gandy and Dekosky, 2012). Diploidy for Apo E4 is believed to contribute to: (i). the aggregation of the toxic neuropeptide amyloid-beta, which can aggregate extracellularly to form insoluble aggregates of amyloid plaques that are a central feature of the Alzheimer’s disease brain; and (ii). defects in clearance of amyloid-beta from the brain (Louveau *et al*., 2016). This association is bolstered by the brain pathologies evident in Down syndrome patients who have extra-copies of chromosome 21 on which the amyloid precursor protein (APP) is located; trisomy results in overexpression of APP (Mentis, 2016). Over two thirds of older adults with trisomy 21 die from dementia, and among them, the risk of premature death is increased by almost seven-fold in APO E4 isoform carriers (Hithersay *et al*., 2018).

AD has been associated with two proteinopathies: amyloid-beta-related pathology and tauopathies. It is now well established that Apo E4 increases levels of amyloid-beta in the brain. Recently, Shi et al (Shi *et al*., 2017) showed that in a mouse model of tauopathy, ApoE e4 also increased the extent of tau-mediated neurodegeneration with increases in brain atrophy and neuroinflammation that appeared to be independent of amyloid-beta pathology. Interestingly, the APO E4 isoform’s principal mechanisms of action seem not to be a function of defects in lipid metabolism pathways, which is the primary activity of APOE in peripheral systems, but rather of effects on a combination of systems implicated in AD. Shi and Holtzman reported that Apo E4 influences immunomodulatory functions of innate immunity, possibly acting through the TREM2 receptor which is expressed on myeloid 2 cells in the CNS microglia (Shi and Holtzman, 2018).

Patients with sporadic AD exhibit impairments in amyloid-beta clearance without changes in the *de novo* production of amyloid-beta (Louveau *et al*., 2016). Indeed, Apo E4 has been shown to directly disrupt clearance of amyloid-beta across the Blood-Brain Barrier (BBB) of mice, providing an explanation for the accumulation of amyloid-beta associated with the APO E4 genotype (Deane *et al*., 2008), and suggesting that impairment in the neurovascular function of the BBB may play a role in AD etiology. Liu et al (Liu *et al*., 2017) developed a cell-type specific, inducible mouse model to control the expression of astrocytic Apo E4 to determine the stage at which it exerts the strongest effect on amyloid pathology. Their data suggests that Apo E4 acts at the seeding stage of amyloid plaque formation, probably by impeding amyloid beta clearance and promoting its aggregation (Liu *et al*., 2017).

## 4. APO E4 in AD-associated neurovascular function

The neurovascular component of AD causality, and more broadly, cognitive dysfunction, has become a field of intense study, inspiring recent calls for further investigation and incorporation of vascular markers into AD research (Sweeney *et al*., 2019). Since amyloid-beta accumulation is associated with cognitive dysfunction, a major point of connection is the neurovascular system’s role in clearing amyloid-beta from the brain. Nation et al. (Nation *et al*., 2019) recently suggested that the disturbance of BBB can, independent of amyloid-beta and tau pathologies, serve as an early biomarker of human cognitive dysfunction, as detected by hippocampal brain capillary damage and BBB breakdown present in individuals with early cognitive dysfunction.

APO E4 is linked to breakdown of the BBB. Bell et al. (Bell *et al*., 2012) showed that in mice, expression of APO E4, but not APO E2 or APO E3 or deficiency of murine apoe4, leads to the breakdown of the BBB as mediated by the vascular pericyte-localized proinflammatory CypA–nuclear factor-kB–matrix-metalloproteinase-9. BBB breakdown leads to neuronal uptake of neurotoxic proteins in the blood and a decrease in the blood flow of the microvasculature and cerebellum. These events occur prior to neuronal dysfunction and may be responsible for the observed neurodegenerative changes (Bell *et al*., 2012).

Cardiovascular risk factors also increase the risk of dementia. Robert et al (Robert *et al*., 2017) used bioengineered human cerebral vessels to show that in that system, Apo E and circulating HDL mediate amyloid-beta transport and Apo E4 was less effective than Apo E2 in facilitating this transport.

## 5. APO E4 and the lymphatic system

Interestingly, brain tissue samples from AD patients show a number of microvascular alterations compared to normal brains, with Bolayannis and Baloyannis reporting *“fusiform dilations, tortuosities, and abnormal branching”* with an overall decrease in the density of capillaries (Baloyannis and Baloyannis, 2012). This study also noted mitochondrial abnormalities in capillary endothelial cells and degeneration of pericytes (Baloyannis and Baloyannis, 2012). The extent to which the lymphatic system of AD patients show these types of abnormalities and Apo E4 affects brain lymphatic vessels is worth investigating. Understanding the branching morphology of these vessels to detect any pathological remodeling of lymphatic vessels in Apo E4-mediated AD could reveal an intriguing new pathologic mechanism.

To this end, Lim et al. (Lim et al., 2009) reported that apoE-deficient (apoE-/-) mice exhibit a number of lymphatic phenotypes, including tissue swelling, leaky lymphatic vessels, a significant dilation of capillary lymphatic vessels, and a reduction in the transport of lymphatic fluid and dendritic cells from peripheral tissue. Moreover, the collecting lymphatic vessels reduce their recruitment of smooth muscle cells and show altered distribution of the lymphatic endothelial hyaluronic acid receptor-1 (LYVE-1) (Lim *et al*., 2009).

It is true, though, that the peripheral and meningeal lymphatic vessels differ in that meningeal lymphatic vessels are less complex with less lymphatic branching and fewer valves to prevent back-flow of lymph fluid (Aspelund *et al*., 2015). Also, interestingly, the metabolic pathways mediating cholesterol homeostasis in the brain also differ from those of peripheral tissues. The BBB prevents peripheral cholesterol from entering the brain and cholesterol is largely synthesized by astrocytes and oligodendrocytes in the brain (Broce *et al*., 2018). However, as discussed above, APO E4 exerts its function on the brain through non-lipid metabolism pathways, notably immunomodulation, whose role in Alzheimer mouse models is increasingly being recognized (Gjoneska *et al*., 2015).

In parallel, endothelial and lymphatic vessels appear to share a close ontogenetic relationship. They share common embryonic cellular origins and lymphatic vessels reprogrammed to become blood vessels affect blood flow-related events such as shear stress (Chen *et al*., 2012). Given the close ontogenetic similarities between capillaries and lymphatics and APO E4’s established role in peripheral lymphatic vessels ((Lim *et al*., 2009) and (Baloyannis and Baloyannis, 2012)), it would be intriguing to know if: (i). meningeal lymphatic vessels play a role in amyloid clearance; and (ii). the extent to which amyloid clearance is compromised by the APO E4 isoform, analogous to its effect on the BBB and glymphatic disruption (Achariyar *et al*., 2016) (Ma *et al*., 2018) – a topic far from being elucidated, notwithstanding the importance of APO E4 in AD.

Da Mesquita et al. recently published two groundbreaking studies showing that ablation of the meningeal lymph vessels in the 5xFAD mouse model of AD results in a striking deposition of amyloid-beta in the meninges, macrophage recruitment to large amyloid-beta aggregates, and an increase in the amyloid-beta plaque load in the hippocampus (Figure 1) (Da Mesquita *et al*., 2018a; Da Mesquita *et al*., 2018b). Moreover, staining for beta-amyloid in the brains of patients with AD and controls without AD revealed a vascular beta-amyloid pathology in the cortical leptomeninges and beta-amyloid depositions in the dura mater adjacent to the superior sagittal sinus (Da Mesquita *et al*., 2018a; Da Mesquita *et al*., 2018b). Together, these findings highlight the importance of meningeal drainage through lymphatic vessels for normal brain physiology and cognitive function. In addition, the worsening of amyloid-beta pathology upon disruption of the meningeal lymphatic system in AD mouse models suggests that dysfunction of the meningeal lymphatic vessels may play an exacerbating role in AD pathology. The AD transgenic mouse models (J20 and 5xFAD) used in the Da Mesquita et al study (Da Mesquita *et al*., 2018b) showed no differences from controls in their meningeal lymphatics.

**Figure 1.**
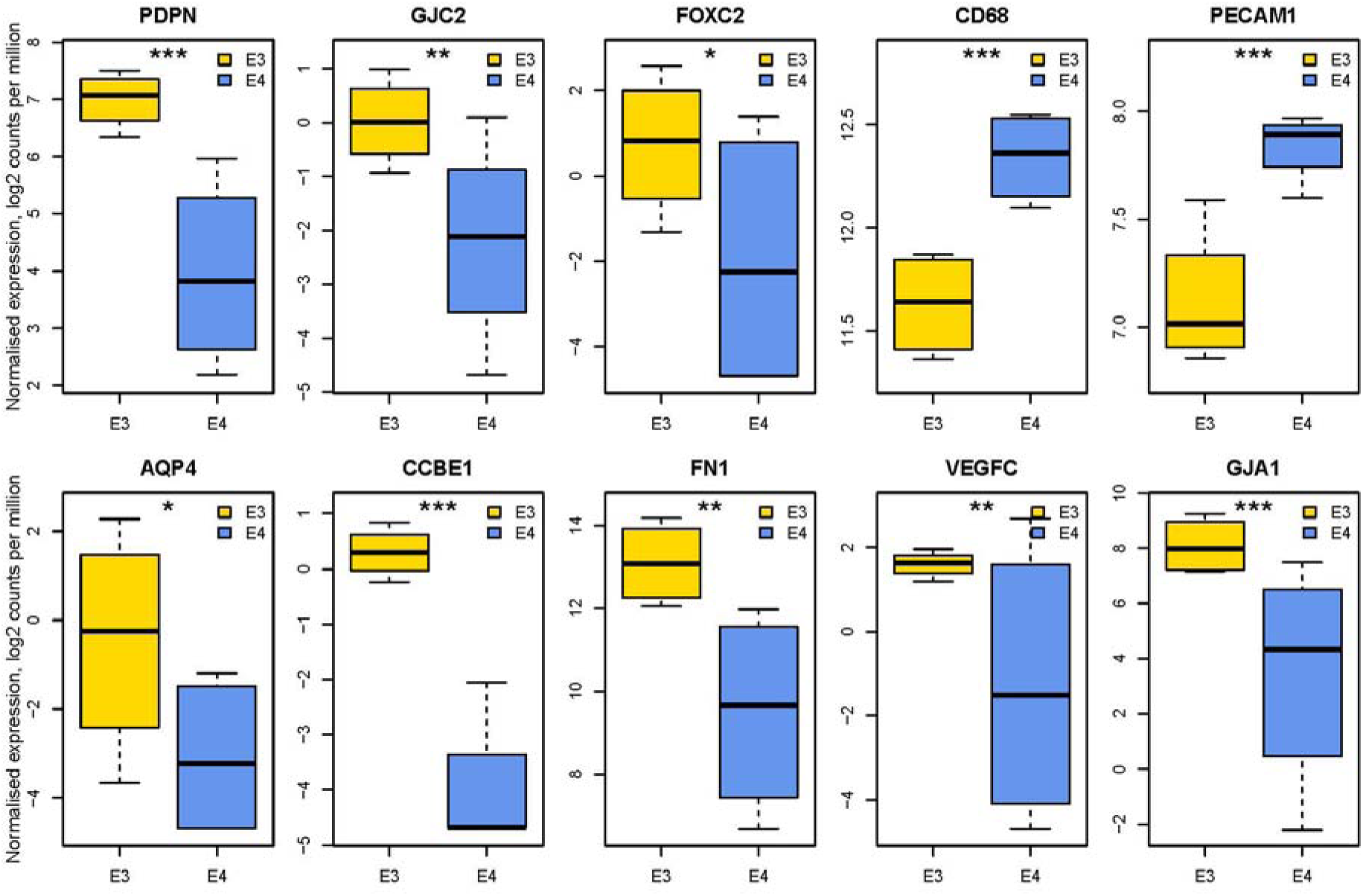
Box and whisker plot showing lymphatic marker expression levels in iPSC-derived neural lineages from APO E4 and APO E3 (control) knock-in cells. Normalized gene expression of lymphatic markers and related genes in cells expressing knock-in of either APO E3 (blue) or APO E4 (yellow). Differentially expressed genes are marked with an asterisk to indicate statistical significance: * for false discovery rate (FDR) <= 0.05; ** for FDR <= 0.01; and *** for FDR <= 0.001. The error bars represent the standard deviation (SD).

However, these mouse models are probably not relevant to APO E4-induced AD, given that they either driven by overexpression of human APP (J20 model) or mutant APP and PSEN expressed in high level (5xFAD model). Hence, they may be more relevant to early-onset AD, while APO E4 more relevant to late-onset AD, and as such, these models may not capture the effects of this major AD risk factor on meningeal lymphatics vessels.

## 6. APO E4, Aquaporin 4, and the meningeal lymphatic and glymphatic systems

In addition to the meningeal lymphatic system, the brain contains the glymphatic system, another system of drainage channels that is involved in the exchange of CSF with ISF and helps clear interstitial waste from the brain parenchyma (the glymphatic system) (Iliff *et al*., 2012). This pathway also was shown to exist in humans (Eide and Ringstad, 2015). When meningeal lymphatic drainage was prevented in mice by the ligation of deep cervical lymphatic nodes (LdcLNs), the disruption of aquaporin 4 (AQP4) water channel, which regulates the glymphatic clearance, resulted in the accumulation of amyloid beta in the hippocampus (Cao *et al*., 2018). The mice with deficits in both glymphatic and meningeal lymphatic clearance exhibited microglial reactivity and activation of the inflammasome, as well as hippocampal neural apoptosis and reduction of cognitive function (Cao *et al*., 2018). Moreover, tau levels were increased in the LdcLNs mice but not the AQP4-null mice (Cao *et al*., 2018). Interestingly, Achariyar et al. (Achariyar *et al*., 2016) showed that the glymphatic system contributes to the transport of lentiviral-delivered Apo E3 to neurons. The authors suggest that, in addition to its role as a clearance system for the brain, the glymphatic systems helps distribute essential molecules through the brain (Achariyar *et al*., 2016).

## 7. Expression of lymphatic-vessel genes in APOE4-expressing cell types of the brain

The cell-specific effects of APO E4 in AD pathology are poorly understood. To determine which human brain cell types are affected by expression of the APO E4 variant, Lin et al examined the effects of CRISPR-mediated APO E4 isogenic homozygous knock-in into human IPSC cells derived from a subject without AD. Using a differentiation system for IPSCs, they were able to generate various brain cell types and compare gene expression resulting from the APO E4 knock-in to that of an analogous APO E3 knock-in. They found that the expression of hundreds of genes was altered in IPSC-derived neurons, astrocytes, and microglia, many of which were also aberrantly expressed in post-mortem samples of AD patients (Lin *et al*., 2018). The observed cellular defects caused by APO E4 expression included: (i). increased synapse number and elevated Aβ_42_ secretion in neurons; (ii). defects in amyloid-beta uptake and cholesterol accumulation in astrocytes; as well as (iii). aberrant morphology correlating with reduced amyloid-beta phagocytosis in microglia (Lin *et al*., 2018).

The dataset affords an opportunity to test the hypothesis that meningeal lymphatic vessels are also affected by APO E4 at the level of gene expression. Therefore, and in light of recent calls to make re-use of open public genomic data (Amann *et al*., 2019), we re-analyzed the cell-type specific RNA-seq data from the Lin et al. study that had been deposited in GEO to identify genes that were differentially expressed in the APO E4 (“pathological” status) knock-in *vs.* APO E3 (control) knock in cells in the starting iPSC, and IPSC-derived neurons, astrocytes, and microglia. We also monitored expression of markers of lymphatic-vessel-related genes, as listed in Table 2.

**Table 2.** Gene markers of human lymphatic vessels.

| **Meningeal lymphatics** | **Initiation of valve formation** |
| --- | --- |
| Prox1 | GATA-2 |
| CD31 (PECAM1) | FOXC2 |
| LYVE-1 | Ephrin type B2 |
| Podoplanin (PDPN) |  |
| VEGFR3 |  |
| CCL21 |  |
| CD68 negativity |  |
| CD45 |  |
| SOS |  |
| **Valve Maturation** | **Lymphedema** |
| Integrin alpha-9 | VEGF-C# |
| Fibronectin 1 (FN 1) | VEGFR-3* |
| Connexin 43 (GJA1) | FOXC2□ |
|  | GATA2* |
|  | CCBE1* |
|  | GJC2* |

\* Mutant gene forms, with functional, missense mutations linked to lymphedema; # VEGFC exposure increases the diameter of meningeal lymphatic vessels, which completely fail to develop upon VEGFC and VEGFD inhibition; □ Marker of initiation of valve formation, and mutant gene forms linked to lymphedema-distichiasis syndrome. Source: Gene markers based on previous reports from: (Louveau *et al.*, 2016); (Louveau *et al.*, 2015) and (2019) OMIM (Online Mendelian Inheritance for Man) Database (www.omim.org) (Assessed January 2019).

Our analysis did not reveal statistically significant differences in the expression of lymphatic vessel genes in iPSC, neurons, and astrocytes (data not shown), based on searches on all the above gene markers. Strikingly, though, the microglial cells showed statistically significant differences between the APO E3 and APO E4 knock-in cells in the expression of several genes specifically related to lymphatics (Fig. 1 and 2). In the APO E3 knock-in cells, we detected increased levels of expression of PDPN, CCBE1, GJA1, FN1, VEGFC, FOXC2, and GJC2, compared to the APO E4 knock-in cells; AQP4 levels were also increased. In the APO E4 knock-in cells types, we detected significant increases in the levels of PECAM1 (CD31) and CD68 (for which absence of “staining” (so-called, CD68 negativity) is linked to lymphatic vessels), compared to APO E3 knock-in cells. Thus, with the exception of CD31, the higher PDPN and lower CD68 levels in APO E3 lines could imply the presence of more pronounced lymphatic markers in APO E3 (see: Figures 1 and 2).

**Figure 2.**
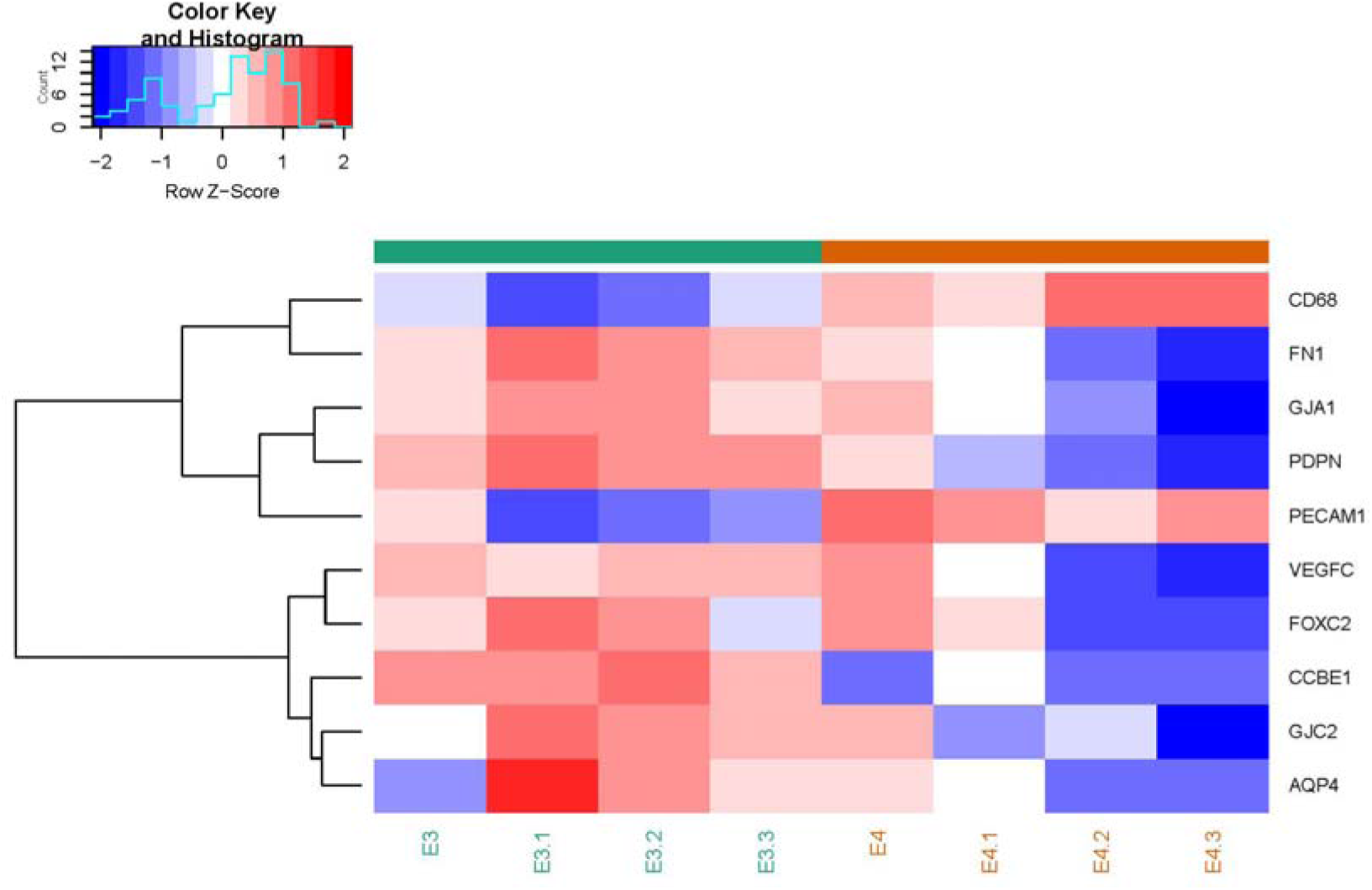
Heat map showing lymphatic marker expression levels in iPSC-derived neural lineages from APO E4 and APO E3 (control) knock-in cells. Samples (x-axis) and genes (y-axis, right) within the heatmap are arranged through hierarchical clustering based on normalized expression values. The heatmap colour is scaled at the gene level to fully visualize the range of expression of each gene.

**Figure 3.**
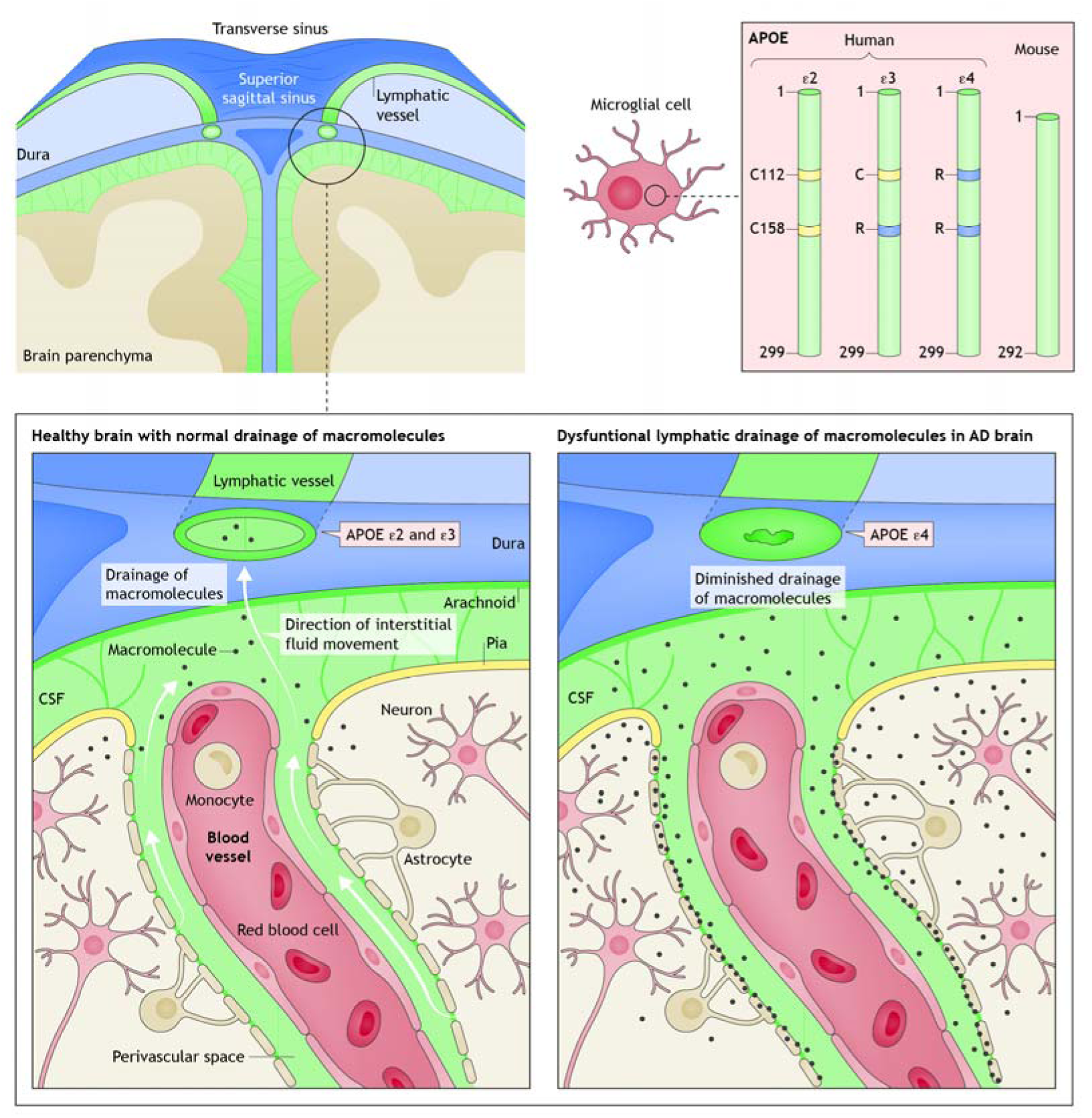
Hypothesized role of Apolipoprotein E (Apo E) and lymphatic vessels in Alzheimer’s disease (AD). Schematic depiction of human and mouse Apo E isoforms (top right), with yellow indicating the presence of cysteine [C] and blue indicating the presence of arginine [R]. Illustration of brain and meninges anatomy in a non-AD brain, showing meningeal lymphatic vessels, brain cell types, and outflow of CSF (top left and magnified in bottom left). The labeling “APO E4” with an arrow pointing to the dysfunctional lymphatic vessel (bottom right) illustrates the hypothesis that APO E4 is associated with the reduction in size of lymphatic vessels and the consequent blockade of clearance of macromolecules such as amyloid-beta. (adapted and modified from Ref. (Gotz *et al*., 2018) © 2018, and ref. (Louveau *et al*., 2015), © 2015, both with permission from Springer Nature)

The microglia cells are ontogenetically related to blood-producing bone marrow cells, and a recent lineage tracing study has shown that their embryonic form (hemogenic endothelial cells) are precursors to lymphatic vessels (Klotz *et al*., 2015). Therefore, a logical extension of the results is that the APOE e3-expressing cell types express more characteristics of lymphatic cells than their APOE e4-expressing counterparts ─ a characteristic of APO E4 which we propose to call “attenuated lymphaticness”. In addition, the lower levels of AQP4 expression in the APO E4 cells could perhaps indicate a disruption in the glymphatic system. Moreover, the results also suggest that APO E4 expression may cause cells to lose their lymphatic characteristics, although *in vitro* studies such as these are limited in their ability to predict true anatomical effects. Furthermore, in APO E4, we observed reduced expression of genes-markers in which functional missense mutations are linked to lymphedema, such as VEGFC (Gordon *et al*., 2013), CCBE1 (Alders *et al*., 2009), FOXC2 (van Steensel *et al*., 2009), and GJC2 (Ferrell *et al*., 2010).

In so doing, these notions suggest that AD, not least in its APOE e4 status, may be linked to shrinkage of meningeal lymphatic vessels and as corollary, reduced lymphatic flow. In other words, if lymphatic vessels undergo shrinkage (“lymphosclerosis”), then the interstitial and CSF fluid will become obstructed in terms of its flow, causing its stagnation (“lymphedema”); however, detailed anatomical studies are essential before deriving any conclusion.

Of note, this “informed hypothesis” may not be entirely surprising, considering that (i). Apo E4 may exercise its effects in a vascular endothelia growth factor (VEGF)-dependent pathway and VEGF upregulation can reverse APO E4 pathology (Salomon-Zimri *et al*., 2016); and (ii). VEGF-C administration can restore meningeal lymphatic vessels pathology in aged mice (Da Mesquita *et al*., 2018b). Nevertheless, APO E4 is the principal genetic factor for AD, so its undiluted mechanistic elucidation remains pivotal, similarly to studies of chief risk factors in other diseases (e.g., mechanistic investigations on FTO region, which has the strongest genetic links to obesity in (Claussnitzer *et al*., 2015)). In addition, a question could be raised: why APO E4 is not implicated in, or already linked to peripheral edema? Based on the Human Protein Atlas (Thul *et al*., 2017), the Apoliprotein E seems to be predominantly expressed at high levels in tissues of cerebral cortex, hippocampus, caudate, and adrenal gland; all of these tissues (considering, of course, only adrenal medulla and not cortex) share common embryological origins (neural tube, and neural crest). Therefore, its chief downstream effects will most likely be in those tissues.

As reviewed by Louveau et al. (Louveau *et al*., 2016), it has remained unclear how meningeal lymphatic vessels respond to and control the high levels of amyloid-beta contained in the brain fluids of AD patients. Possibilities include drainage into the CSF, paravascular clearance, clearance through the glymphatic pathway, or other mechanisms (Louveau *et al*., 2016). Given that disruption of meningeal lymphatic vessels in AD mouse model leads to amyloid-beta deposits in the meninges, these questions take on new urgency (Da Mesquita *et al*., 2018b). We suggest that future studies should seek to clarify the extent to which and mechanisms whereby APO E4 influences amyloid-beta deposition in the meninges. This issue may be clarified by electron microscopy or cryo-electron microscopy of APO E4 expressing cells to elucidate abnormalities in the morphology of vasculature and lymphatic system, notably the diameter of lymphatic vessels and their branching morphology. The 7-Tesla MRI imaging approach of Absinta et al. (Absinta *et al*., 2017), who first described the meningeal lymphatic vessels in humans, could enable measurements of their diameters in examinations of AD patients who were carriers or non-carriers of the APOE e4 variant (Absinta *et al*., 2017).

## 8. Conclusions

Clearly, the field of meningeal lymphatics is ripe for investigation. The recent studies discussed herein have raised fascinating questions about the connection between APO E4, the meningeal lymphatic vessels, and AD. Deciphering the role of APO 4, which so far is the strongest genetic link to AD, in the brain’s lymphatic system could reveal a missing link in our understanding of the etiology and pathology of AD. Hence, we hope this perspective coupled with previous studies’ findings may help in formulating the concept of APO E4-related AD (such as in late onset AD or trisomy 21-mediated AD) lymphosclerosis coupled with lymphedema. In addition, our approach highlights the power of open data and re-analysis to offer new perspectives, in alignment with recent calls ((Amann *et al*., 2019)).

## 9. Appendix: Differential Expression (re)analysis

Processed gene-count matrices for each of the cell lineages were obtained from the GEO repository (accession number GSE102956) based on (Lin *et al*., 2018). Counts were merged into a single expression matrix and genes with average log 2 counts per million (CPM) across all samples that fell below −0.75 were removed from the analysis to avoid inflated dispersion affecting genes with low expression. This filter removed 5342 genes leaving 15382 genes for analysis. Normalization factors were calculated using the “Trimmed mean of M values (TMM)” method (Robinson and Oshlack, 2010) in edgeR package (Robinson *et al*., 2010). These normalization factors were used to calculate effective library sizes, which in turn, were employed to calculate normalized CPM values for genes of interest and visualized as boxplots and a heatmap. We used the voom method (Law *et al*., 2014) to transform counts and derive gene-level weights. The transformed data was used to model gene expression jointly across all cell types and experimental conditions using a linear model framework in limma (Ritchie *et al*., 2015). Raw p values were adjusted for multiple testing by applying the Benjamini-Hochberg method (Benjamini and Hochberg, 1995) on the 20 genes of interest.

## References

1. (2019) Online Mendelian Inheritance in Man. Ed. J.H. University. Johns Hopkins University

2. Absinta, M., Ha, S.K., Nair, G., Sati, P., Luciano, N.J., Palisoc, M., Louveau, A., Zaghloul, K.A., Pittaluga, S., Kipnis, J., Reich, D.S., 2017. Human and nonhuman primate meninges harbor lymphatic vessels that can be visualized noninvasively by MRI. eLife 6.

3. Achariyar, T.M., Li, B., Peng, W., Verghese, P.B., Shi, Y., McConnell, E., Benraiss, A., Kasper, T., Song, W., Takano, T., Holtzman, D.M., Nedergaard, M., Deane, R., 2016. Glymphatic distribution of CSF-derived apoE into brain is isoform specific and suppressed during sleep deprivation. Molecular neurodegeneration 11, 74.

4. Alders, M., Al-Gazali, L., Cordeiro, I., Dallapiccola, B., Garavelli, L., Tuysuz, B., Salehi, F., Haagmans, M.A., Mook, O.R., Majoie, C.B., Mannens, M.M., Hennekam, R.C., 2014. Hennekam syndrome can be caused by FAT4 mutations and be allelic to Van Maldergem syndrome. Human genetics 133, 1161–1167.

5. Alders, M., Hogan, B.M., Gjini, E., Salehi, F., Al-Gazali, L., Hennekam, E.A., Holmberg, E.E., Mannens, M.M., Mulder, M.F., Offerhaus, G.J., Prescott, T.E., Schroor, E.J., Verheij, J.B., Witte, M., Zwijnenburg, P.J., Vikkula, M., Schulte-Merker, S., Hennekam, R.C., 2009. Mutations in CCBE1 cause generalized lymph vessel dysplasia in humans. Nature genetics 41, 1272–1274.

6. Amann, R.I., Baichoo, S., Blencowe, B.J., Bork, P., Borodovsky, M., Brooksbank, C., Chain, P.S.G., Colwell, R.R., Daffonchio, D.G., Danchin, A., de Lorenzo, V., Dorrestein, P.C., Finn, R.D., Fraser, C.M., Gilbert, J.A., Hallam, S.J., Hugenholtz, P., Ioannidis, J.P.A., Jansson, J.K., Kim, J.F., Klenk, H.P., Klotz, M.G., Knight, R., Konstantinidis, K.T., Kyrpides, N.C., Mason, C.E., McHardy, A.C., Meyer, F., Ouzounis, C.A., Patrinos, A.A.N., Podar, M., Pollard, K.S., Ravel, J., Munoz, A.R., Roberts, R.J., Rossello-Mora, R., Sansone, S.A., Schloss, P.D., Schriml, L.M., Setubal, J.C., Sorek, R., Stevens, R.L., Tiedje, J.M., Turjanski, A., Tyson, G.W., Ussery, D.W., Weinstock, G.M., White, O., Whitman, W.B., Xenarios, I., 2019. Toward unrestricted use of public genomic data. Science (New York, N.Y.) 363, 350–352.

7. Andres, K.H., von During, M., Muszynski, K., Schmidt, R.F., 1987. Nerve fibres and their terminals of the dura mater encephali of the rat. Anatomy and embryology 175, 289–301.

8. Antila, S., Karaman, S., Nurmi, H., Airavaara, M., Voutilainen, M.H., Mathivet, T., Chilov, D., Li, Z., Koppinen, T., Park, J.H., Fang, S., Aspelund, A., Saarma, M., Eichmann, A., Thomas, J.L., Alitalo, K., 2017. Development and plasticity of meningeal lymphatic vessels. The Journal of experimental medicine 214, 3645–3667.

9. Aspelund, A., Antila, S., Proulx, S.T., Karlsen, T.V., Karaman, S., Detmar, M., Wiig, H., Alitalo, K., 2015. A dural lymphatic vascular system that drains brain interstitial fluid and macromolecules. The Journal of experimental medicine 212, 991–999.

10. Baloyannis, S.J., Baloyannis, I.S., 2012. The vascular factor in Alzheimer’s disease: a study in Golgi technique and electron microscopy. Journal of the neurological sciences 322, 117–121.

11. Bell, R.D., Winkler, E.A., Singh, I., Sagare, A.P., Deane, R., Wu, Z., Holtzman, D.M., Betsholtz, C., Armulik, A., Sallstrom, J., Berk, B.C., Zlokovic, B.V., 2012. Apolipoprotein E controls cerebrovascular integrity via cyclophilin A. Nature 485, 512–516.

12. Bellou, V., Belbasis, L., Tzoulaki, I., Middleton, L.T., Ioannidis, J.P.A., Evangelou, E., 2017. Systematic evaluation of the associations between environmental risk factors and dementia: An umbrella review of systematic reviews and meta-analyses. Alzheimer’s & dementia: the journal of the Alzheimer’s Association 13, 406–418.

13. Benjamini, Y., Hochberg, Y., 1995. Controlling the false discovery rate: a practical and powerful approach to multiple testing. Journal of the Royal statistical society: series B (Methodological) 57, 289–300.

14. Berton, M., Lorette, G., Baulieu, F., Lagrue, E., Blesson, S., Cambazard, F., Vaillant, L., Maruani, A., 2015. Generalized lymphedema associated with neurologic signs (GLANS) syndrome: a new entity? Journal of the American Academy of Dermatology 72, 333–339.

15. Broce, I.J., Tan, C.H., Fan, C.C., Jansen, I., Savage, J.E., Witoelar, A., Wen, N., Hess, C.P., Dillon, W.P., Glastonbury, C.M., Glymour, M., Yokoyama, J.S., Elahi, F.M., Rabinovici, G.D., Miller, B.L., Mormino, E.C., Sperling, R.A., Bennett, D.A., McEvoy, L.K., Brewer, J.B., Feldman, H.H., Hyman, B.T., Pericak-Vance, M., Haines, J.L., Farrer, L.A., Mayeux, R., Schellenberg, G.D., Yaffe, K., Sugrue, L.P., Dale, A.M., Posthuma, D., Andreassen, O.A., Karch, C.M., Desikan, R.S., 2018. Dissecting the genetic relationship between cardiovascular risk factors and Alzheimer’s disease. Acta neuropathologica.

16. Cai, R., Pan, C., Ghasemigharagoz, A., Todorov, M.I., Forstera, B., Zhao, S., Bhatia, H.S., Parra-Damas, A., Mrowka, L., Theodorou, D., Rempfler, M., Xavier, A.L.R., Kress, B.T., Benakis, C., Steinke, H., Liebscher, S., Bechmann, I., Liesz, A., Menze, B., Kerschensteiner, M., Nedergaard, M., Erturk, A., 2019. Panoptic imaging of transparent mice reveals whole-body neuronal projections and skull-meninges connections. Nature neuroscience 22, 317–327.

17. Cao, X., Xu, H., Feng, W., Su, D., Xiao, M., 2018. Deletion of aquaporin-4 aggravates brain pathology after blocking of the meningeal lymphatic drainage. Brain research bulletin 143, 83–96.

18. Chen, C.Y., Bertozzi, C., Zou, Z., Yuan, L., Lee, J.S., Lu, M., Stachelek, S.J., Srinivasan, S., Guo, L., Vicente, A., Mericko, P., Levy, R.J., Makinen, T., Oliver, G., Kahn, M.L., 2012. Blood flow reprograms lymphatic vessels to blood vessels. The Journal of clinical investigation 122, 2006–2017.

19. Chrousos, G.P., 1995. The hypothalamic-pituitary-adrenal axis and immune-mediated inflammation. The New England journal of medicine 332, 1351–1362.

20. Claussnitzer, M., Dankel, S.N., Kim, K.H., Quon, G., Meuleman, W., Haugen, C., Glunk, V., Sousa, I.S., Beaudry, J.L., Puviindran, V., Abdennur, N.A., Liu, J., Svensson, P.A., Hsu, Y.H., Drucker, D.J., Mellgren, G., Hui, C.C., Hauner, H., Kellis, M., 2015. FTO Obesity Variant Circuitry and Adipocyte Browning in Humans. The New England journal of medicine 373, 895–907.

21. Da Mesquita, S., Fu, Z., Kipnis, J., 2018a. The Meningeal Lymphatic System: A New Player in Neurophysiology. Neuron 100, 375–388.

22. Da Mesquita, S., Louveau, A., Vaccari, A., Smirnov, I., Cornelison, R.C., Kingsmore, K.M., Contarino, C., Onengut-Gumuscu, S., Farber, E., Raper, D., Viar, K.E., Powell, R.D., Baker, W., Dabhi, N., Bai, R., Cao, R., Hu, S., Rich, S.S., Munson, J.M., Lopes, M.B., Overall, C.C., Acton, S.T., Kipnis, J., 2018b. Functional aspects of meningeal lymphatics in ageing and Alzheimer’s disease. Nature 560, 185–191.

23. Deane, R., Sagare, A., Hamm, K., Parisi, M., Lane, S., Finn, M.B., Holtzman, D.M., Zlokovic, B.V., 2008. apoE isoform-specific disruption of amyloid beta peptide clearance from mouse brain. The Journal of clinical investigation 118, 4002–4013.

24. Eide, P.K., Ringstad, G., 2015. MRI with intrathecal MRI gadolinium contrast medium administration: a possible method to assess glymphatic function in human brain. Acta radiologica open 4, 2058460115609635.

25. Ferrell, R.E., Baty, C.J., Kimak, M.A., Karlsson, J.M., Lawrence, E.C., Franke-Snyder, M., Meriney, S.D., Feingold, E., Finegold, D.N., 2010. GJC2 missense mutations cause human lymphedema. American journal of human genetics 86, 943–948.

26. Foldi, M., Gellert, A., Kozma, M., Poberai, M., Zoltan, O.T., Csanda, E., 1966. New contributions to the anatomical connections of the brain and the lymphatic system. Acta anatomica 64, 498–505.

27. Furukawa, M., Shimoda, H., Kajiwara, T., Kato, S., Yanagisawa, S., 2008. Topographic study on nerve-associated lymphatic vessels in the murine craniofacial region by immunohistochemistry and electron microscopy. Biomedical research (Tokyo, Japan) 29, 289–296.

28. Gandy, S., Dekosky, S.T., 2012. APOE epsilon4 status and traumatic brain injury on the gridiron or the battlefield. Science translational medicine 4, 134ed134.

29. Gassen, N.C., Chrousos, G.P., Binder, E.B., Zannas, A.S., 2017. Life stress, glucocorticoid signaling, and the aging epigenome: Implications for aging-related diseases. Neuroscience and biobehavioral reviews 74, 356–365.

30. Gausas, R.E., Daly, T., Fogt, F., 2007. D2-40 expression demonstrates lymphatic vessel characteristics in the dural portion of the optic nerve sheath. Ophthalmic plastic and reconstructive surgery 23, 32–36.

31. Gjoneska, E., Pfenning, A.R., Mathys, H., Quon, G., Kundaje, A., Tsai, L.H., Kellis, M., 2015. Conserved epigenomic signals in mice and humans reveal immune basis of Alzheimer’s disease. Nature 518, 365–369.

32. Gordon, K., Schulte, D., Brice, G., Simpson, M.A., Roukens, M.G., van Impel, A., Connell, F., Kalidas, K., Jeffery, S., Mortimer, P.S., Mansour, S., Schulte-Merker, S., Ostergaard, P., 2013. Mutation in vascular endothelial growth factor-C, a ligand for vascular endothelial growth factor receptor-3, is associated with autosomal dominant milroy-like primary lymphedema. Circulation research 112, 956–960.

33. Gotz, J., Bodea, L.G., Goedert, M., 2018. Rodent models for Alzheimer disease. Nature reviews. Neuroscience 19, 583–598.

34. Hithersay, R., Startin, C.M., Hamburg, S., Mok, K.Y., Hardy, J., Fisher, E.M.C., Tybulewicz, V.L.J., Nizetic, D., Strydom, A., 2018. Association of Dementia With Mortality Among Adults With Down Syndrome Older Than 35 Years. JAMA neurology.

35. Iliff, J.J., Wang, M., Liao, Y., Plogg, B.A., Peng, W., Gundersen, G.A., Benveniste, H., Vates, G.E., Deane, R., Goldman, S.A., Nagelhus, E.A., Nedergaard, M., 2012. A paravascular pathway facilitates CSF flow through the brain parenchyma and the clearance of interstitial solutes, including amyloid beta. Science translational medicine 4, 147ra111.

36. Jansen, I.E., Savage, J.E., Watanabe, K., Bryois, J., Williams, D.M., Steinberg, S., Sealock, J., Karlsson, I.K., Hagg, S., Athanasiu, L., Voyle, N., Proitsi, P., Witoelar, A., Stringer, S., Aarsland, D., Almdahl, I.S., Andersen, F., Bergh, S., Bettella, F., Bjornsson, S., Braekhus, A., Brathen, G., de Leeuw, C., Desikan, R.S., Djurovic, S., Dumitrescu, L., Fladby, T., Hohman, T.J., Jonsson, P.V., Kiddle, S.J., Rongve, A., Saltvedt, I., Sando, S.B., Selbaek, G., Shoai, M., Skene, N.G., Snaedal, J., Stordal, E., Ulstein, I.D., Wang, Y., White, L.R., Hardy, J., Hjerling-Leffler, J., Sullivan, P.F., van der Flier, W.M., Dobson, R., Davis, L.K., Stefansson, H., Stefansson, K., Pedersen, N.L., Ripke, S., Andreassen, O.A., Posthuma, D., 2019. Genome-wide meta-analysis identifies new loci and functional pathways influencing Alzheimer’s disease risk. Nature genetics 51, 404–413.

37. Klotz, L., Norman, S., Vieira, J.M., Masters, M., Rohling, M., Dube, K.N., Bollini, S., Matsuzaki, F., Carr, C.A., Riley, P.R., 2015. Cardiac lymphatics are heterogeneous in origin and respond to injury. Nature 522, 62–67.

38. Kunkle, B.W., Grenier-Boley, B., Sims, R., Bis, J.C., Damotte, V., Naj, A.C., Boland, A., Vronskaya, M., van der Lee, S.J., Amlie-Wolf, A., Bellenguez, C., Frizatti, A., Chouraki, V., Martin, E.R., Sleegers, K., Badarinarayan, N., Jakobsdottir, J., Hamilton-Nelson, K.L., Moreno-Grau, S., Olaso, R., Raybould, R., Chen, Y., Kuzma, A.B., Hiltunen, M., Morgan, T., Ahmad, S., Vardarajan, B.N., Epelbaum, J., Hoffmann, P., Boada, M., Beecham, G.W., Garnier, J.G., Harold, D., Fitzpatrick, A.L., Valladares, O., Moutet, M.L., Gerrish, A., Smith, A.V., Qu, L., Bacq, D., Denning, N., Jian, X., Zhao, Y., Del Zompo, M., Fox, N.C., Choi, S.H., Mateo, I., Hughes, J.T., Adams, H.H., Malamon, J., Sanchez-Garcia, F., Patel, Y., Brody, J.A., Dombroski, B.A., Naranjo, M.C.D., Daniilidou, M., Eiriksdottir, G., Mukherjee, S., Wallon, D., Uphill, J., Aspelund, T., Cantwell, L.B., Garzia, F., Galimberti, D., Hofer, E., Butkiewicz, M., Fin, B., Scarpini, E., Sarnowski, C., Bush, W.S., Meslage, S., Kornhuber, J., White, C.C., Song, Y., Barber, R.C., Engelborghs, S., Sordon, S., Voijnovic, D., Adams, P.M., Vandenberghe, R., Mayhaus, M., Cupples, L.A., Albert, M.S., De Deyn, P.P., Gu, W., Himali, J.J., Beekly, D., Squassina, A., Hartmann, A.M., Orellana, A., Blacker, D., Rodriguez-Rodriguez, E., Lovestone, S., Garcia, M.E., Doody, R.S., Munoz-Fernadez, C., Sussams, R., Lin, H., Fairchild, T.J., Benito, Y.A., Holmes, C., Karamujic-Comic, H., Frosch, M.P., Thonberg, H., Maier, W., Roschupkin, G., Ghetti, B., Giedraitis, V., Kawalia, A., Li, S., Huebinger, R.M., Kilander, L., Moebus, S., Hernandez, I., Kamboh, M.I., Brundin, R., Turton, J., Yang, Q., Katz, M.J., Concari, L., Lord, J., Beiser, A.S., Keene, C.D., Helisalmi, S., Kloszewska, I., Kukull, W.A., Koivisto, A.M., Lynch, A., Tarraga, L., Larson, E.B., Haapasalo, A., Lawlor, B., Mosley, T.H., Lipton, R.B., Solfrizzi, V., Gill, M., Longstreth, W.T., Jr., Montine, T.J., Frisardi, V., Diez-Fairen, M., Rivadeneira, F., Petersen, R.C., Deramecourt, V., Alvarez, I., Salani, F., Ciaramella, A., Boerwinkle, E., Reiman, E.M., Fievet, N., Rotter, J.I., Reisch, J.S., Hanon, O., Cupidi, C., Andre Uitterlinden, A.G., Royall, D.R., Dufouil, C., Maletta, R.G., de Rojas, I., Sano, M., Brice, A., Cecchetti, R., George-Hyslop, P.S., Ritchie, K., Tsolaki, M., Tsuang, D.W., Dubois, B., Craig, D., Wu, C.K., Soininen, H., Avramidou, D., Albin, R.L., Fratiglioni, L., Germanou, A., Apostolova, L.G., Keller, L., Koutroumani, M., Arnold, S.E., Panza, F., Gkatzima, O., Asthana, S., Hannequin, D., Whitehead, P., Atwood, C.S., Caffarra, P., Hampel, H., Quintela, I., Carracedo, A., Lannfelt, L., Rubinsztein, D.C., Barnes, L.L., Pasquier, F., Frolich, L., Barral, S., McGuinness, B., Beach, T.G., Johnston, J.A., Becker, J.T., Passmore, P., Bigio, E.H., Schott, J.M., Bird, T.D., Warren, J.D., Boeve, B.F., Lupton, M.K., Bowen, J.D., Proitsi, P., Boxer, A., Powell, J.F., Burke, J.R., Kauwe, J.S.K., Burns, J.M., Mancuso, M., Buxbaum, J.D., Bonuccelli, U., Cairns, N.J., McQuillin, A., Cao, C., Livingston, G., Carlson, C.S., Bass, N.J., Carlsson, C.M., Hardy, J., Carney, R.M., Bras, J., Carrasquillo, M.M., Guerreiro, R., Allen, M., Chui, H.C., Fisher, E., Masullo, C., Crocco, E.A., DeCarli, C., Bisceglio, G., Dick, M., Ma, L., Duara, R., Graff-Radford, N.R., Evans, D.A., Hodges, A., Faber, K.M., Scherer, M., Fallon, K.B., Riemenschneider, M., Fardo, D.W., Heun, R., Farlow, M.R., Kolsch, H., Ferris, S., Leber, M., Foroud, T.M., Heuser, I., Galasko, D.R., Giegling, I., Gearing, M., Hull, M., Geschwind, D.H., Gilbert, J.R., Morris, J., Green, R.C., Mayo, K., Growdon, J.H., Feulner, T., Hamilton, R.L., Harrell, L.E., Drichel, D., Honig, L.S., Cushion, T.D., Huentelman, M.J., Hollingworth, P., Hulette, C.M., Hyman, B.T., Marshall, R., Jarvik, G.P., Meggy, A., Abner, E., Menzies, G.E., Jin, L.W., Leonenko, G., Real, L.M., Jun, G.R., Baldwin, C.T., Grozeva, D., Karydas, A., Russo, G., Kaye, J.A., Kim, R., Jessen, F., Kowall, N.W., Vellas, B., Kramer, J.H., Vardy, E., LaFerla, F.M., Jockel, K.H., Lah, J.J., Dichgans, M., Leverenz, J.B., Mann, D., Levey, A.I., Pickering-Brown, S., Lieberman, A.P., Klopp, N., Lunetta, K.L., Wichmann, H.E., Lyketsos, C.G., Morgan, K., Marson, D.C., Brown, K., Martiniuk, F., Medway, C., Mash, D.C., Nothen, M.M., Masliah, E., Hooper, N.M., McCormick, W.C., Daniele, A., McCurry, S.M., Bayer, A., McDavid, A.N., Gallacher, J., McKee, A.C., van den Bussche, H., Mesulam, M., Brayne, C., Miller, B.L., Riedel-Heller, S., Miller, C.A., Miller, J.W., Al-Chalabi, A., Morris, J.C., Shaw, C.E., Myers, A.J., Wiltfang, J., O’Bryant, S., Olichney, J.M., Alvarez, V., Parisi, J.E., Singleton, A.B., Paulson, H.L., Collinge, J., Perry, W.R., Mead, S., Peskind, E., Cribbs, D.H., Rossor, M., Pierce, A., Ryan, N.S., Poon, W.W., Nacmias, B., Potter, H., Sorbi, S., Quinn, J.F., Sacchinelli, E., Raj, A., Spalletta, G., Raskind, M., Caltagirone, C., Bossu, P., Orfei, M.D., Reisberg, B., Clarke, R., Reitz, C., Smith, A.D., Ringman, J.M., Warden, D., Roberson, E.D., Wilcock, G., Rogaeva, E., Bruni, A.C., Rosen, H.J., Gallo, M., Rosenberg, R.N., Ben-Shlomo, Y., Sager, M.A., Mecocci, P., Saykin, A.J., Pastor, P., Cuccaro, M.L., Vance, J.M., Schneider, J.A., Schneider, L.S., Slifer, S., Seeley, W.W., Smith, A.G., Sonnen, J.A., Spina, S., Stern, R.A., Swerdlow, R.H., Tang, M., Tanzi, R.E., Trojanowski, J.Q., Troncoso, J.C., Van Deerlin, V.M., Van Eldik, L.J., Vinters, H.V., Vonsattel, J.P., Weintraub, S., Welsh-Bohmer, K.A., Wilhelmsen, K.C., Williamson, J., Wingo, T.S., Woltjer, R.L., Wright, C.B., Yu, C.E., Yu, L., Saba, Y., Pilotto, A., Bullido, M.J., Peters, O., Crane, P.K., Bennett, D., Bosco, P., Coto, E., Boccardi, V., De Jager, P.L., Lleo, A., Warner, N., Lopez, O.L., Ingelsson, M., Deloukas, P., Cruchaga, C., Graff, C., Gwilliam, R., Fornage, M., Goate, A.M., Sanchez-Juan, P., Kehoe, P.G., Amin, N., Ertekin-Taner, N., Berr, C., Debette, S., Love, S., Launer, L.J., Younkin, S.G., Dartigues, J.F., Corcoran, C., Ikram, M.A., Dickson, D.W., Nicolas, G., Campion, D., Tschanz, J., Schmidt, H., Hakonarson, H., Clarimon, J., Munger, R., Schmidt, R., Farrer, L.A., Van Broeckhoven, C., M, C.O.D., DeStefano, A.L., Jones, L., Haines, J.L., Deleuze, J.F., Owen, M.J., Gudnason, V., Mayeux, R., Escott-Price, V., Psaty, B.M., Ramirez, A., Wang, L.S., Ruiz, A., van Duijn, C.M., Holmans, P.A., Seshadri, S., Williams, J., Amouyel, P., Schellenberg, G.D., Lambert, J.C., Pericak-Vance, M.A., 2019. Genetic meta-analysis of diagnosed Alzheimer’s disease identifies new risk loci and implicates Abeta, tau, immunity and lipid processing. Nature genetics 51, 414–430.

39. Lanoiselee, H.M., Nicolas, G., Wallon, D., Rovelet-Lecrux, A., Lacour, M., Rousseau, S., Richard, A.C., Pasquier, F., Rollin-Sillaire, A., Martinaud, O., Quillard-Muraine, M., de la Sayette, V., Boutoleau-Bretonniere, C., Etcharry-Bouyx, F., Chauvire, V., Sarazin, M., le Ber, I., Epelbaum, S., Jonveaux, T., Rouaud, O., Ceccaldi, M., Felician, O., Godefroy, O., Formaglio, M., Croisile, B., Auriacombe, S., Chamard, L., Vincent, J.L., Sauvee, M., Marelli-Tosi, C., Gabelle, A., Ozsancak, C., Pariente, J., Paquet, C., Hannequin, D., Campion, D., 2017. APP, PSEN1, and PSEN2 mutations in early-onset Alzheimer disease: A genetic screening study of familial and sporadic cases. PLoS medicine 14, e1002270.

40. Law, C.W., Chen, Y., Shi, W., Smyth, G.K., 2014. voom: Precision weights unlock linear model analysis tools for RNA-seq read counts. Genome biology 15, R29.

41. Le, C.P., Nowell, C.J., Kim-Fuchs, C., Botteri, E., Hiller, J.G., Ismail, H., Pimentel, M.A., Chai, M.G., Karnezis, T., Rotmensz, N., Renne, G., Gandini, S., Pouton, C.W., Ferrari, D., Moller, A., Stacker, S.A., Sloan, E.K., 2016a. Chronic stress in mice remodels lymph vasculature to promote tumour cell dissemination. Nature communications 7, 10634.

42. Le, M.H., Weissmiller, A.M., Monte, L., Lin, P.H., Hexom, T.C., Natera, O., Wu, C., Rissman, R.A., 2016b. Functional Impact of Corticotropin-Releasing Factor Exposure on Tau Phosphorylation and Axon Transport. PloS one 11, e0147250.

43. Lim, H.Y., Rutkowski, J.M., Helft, J., Reddy, S.T., Swartz, M.A., Randolph, G.J., Angeli, V., 2009. Hypercholesterolemic mice exhibit lymphatic vessel dysfunction and degeneration. The American journal of pathology 175, 1328–1337.

44. Lin, Y.T., Seo, J., Gao, F., Feldman, H.M., Wen, H.L., Penney, J., Cam, H.P., Gjoneska, E., Raja, W.K., Cheng, J., Rueda, R., Kritskiy, O., Abdurrob, F., Peng, Z., Milo, B., Yu, C.J., Elmsaouri, S., Dey, D., Ko, T., Yankner, B.A., Tsai, L.H., 2018. APOE4 Causes Widespread Molecular and Cellular Alterations Associated with Alzheimer’s Disease Phenotypes in Human iPSC-Derived Brain Cell Types. Neuron 98, 1141–1154.e1147.

45. Liu, C.C., Zhao, N., Fu, Y., Wang, N., Linares, C., Tsai, C.W., Bu, G., 2017. ApoE4 Accelerates Early Seeding of Amyloid Pathology. Neuron 96, 1024–1032.e1023.

46. Louveau, A., Da Mesquita, S., Kipnis, J., 2016. Lymphatics in Neurological Disorders: A Neuro-Lympho-Vascular Component of Multiple Sclerosis and Alzheimer’s Disease? Neuron 91, 957–973.

47. Louveau, A., Smirnov, I., Keyes, T.J., Eccles, J.D., Rouhani, S.J., Peske, J.D., Derecki, N.C., Castle, D., Mandell, J.W., Lee, K.S., Harris, T.H., Kipnis, J., 2015. Structural and functional features of central nervous system lymphatic vessels. Nature 523, 337–341.

48. Lukic, I.K., Gluncic, V., Ivkic, G., Hubenstorf, M., Marusic, A., 2003. Virtual dissection: a lesson from the 18th century. Lancet (London, England) 362, 2110–2113.

49. Ma, Q., Ineichen, B.V., Detmar, M., Proulx, S.T., 2017. Outflow of cerebrospinal fluid is predominantly through lymphatic vessels and is reduced in aged mice. Nature communications 8, 1434.

50. Ma, Q., Zhao, Z., Sagare, A.P., Wu, Y., Wang, M., Owens, N.C., Verghese, P.B., Herz, J., Holtzman, D.M., Zlokovic, B.V., 2018. Blood-brain barrier-associated pericytes internalize and clear aggregated amyloid-beta42 by LRP1-dependent apolipoprotein E isoform-specific mechanism. Molecular neurodegeneration 13, 57.

51. Martinez-Corral, I., Ulvmar, M.H., Stanczuk, L., Tatin, F., Kizhatil, K., John, S.W., Alitalo, K., Ortega, S., Makinen, T., 2015. Nonvenous origin of dermal lymphatic vasculature. Circulation research 116, 1649–1654.

52. Mentis, A.F., 2016. Epigenomic engineering for Down syndrome. Neuroscience and biobehavioral reviews 71, 323–327.

53. Nation, D.A., Sweeney, M.D., Montagne, A., Sagare, A.P., D’Orazio, L.M., Pachicano, M., Sepehrband, F., Nelson, A.R., Buennagel, D.P., Harrington, M.G., Benzinger, T.L.S., Fagan, A.M., Ringman, J.M., Schneider, L.S., Morris, J.C., Chui, H.C., Law, M., Toga, A.W., Zlokovic, B.V., 2019. Blood-brain barrier breakdown is an early biomarker of human cognitive dysfunction. Nature medicine 25, 270–276.

54. Nicolaides, N.C., Kyratzi, E., Lamprokostopoulou, A., Chrousos, G.P., Charmandari, E., 2015. Stress, the stress system and the role of glucocorticoids. Neuroimmunomodulation 22, 6–19.

55. Pichol-Thievend, C., Betterman, K.L., Liu, X., Ma, W., Skoczylas, R., Lesieur, E., Bos, F.L., Schulte, D., Schulte-Merker, S., Hogan, B.M., Oliver, G., Harvey, N.L., Francois, M., 2018. A blood capillary plexus-derived population of progenitor cells contributes to genesis of the dermal lymphatic vasculature during embryonic development. Development (Cambridge, England) 145.

56. Prineas, J.W., 1979. Multiple sclerosis: presence of lymphatic capillaries and lymphoid tissue in the brain and spinal cord. Science (New York, N.Y.) 203, 1123–1125.

57. Psaltopoulou, T., Sergentanis, T.N., Panagiotakos, D.B., Sergentanis, I.N., Kosti, R., Scarmeas, N., 2013. Mediterranean diet, stroke, cognitive impairment, and depression: A meta-analysis. Annals of neurology 74, 580–591.

58. Ritchie, M.E., Phipson, B., Wu, D., Hu, Y., Law, C.W., Shi, W., Smyth, G.K., 2015. limma powers differential expression analyses for RNA-sequencing and microarray studies. Nucleic acids research 43, e47.

59. Robert, J., Button, E.B., Yuen, B., Gilmour, M., Kang, K., Bahrabadi, A., Stukas, S., Zhao, W., Kulic, I., Wellington, C.L., 2017. Clearance of beta-amyloid is facilitated by apolipoprotein E and circulating high-density lipoproteins in bioengineered human vessels. eLife 6.

60. Robinson, M.D., McCarthy, D.J., Smyth, G.K., 2010. edgeR: a Bioconductor package for differential expression analysis of digital gene expression data. Bioinformatics (Oxford, England) 26, 139–140.

61. Robinson, M.D., Oshlack, A., 2010. A scaling normalization method for differential expression analysis of RNA-seq data. Genome biology 11, R25.

62. Salomon-Zimri, S., Glat, M.J., Barhum, Y., Luz, I., Boehm-Cagan, A., Liraz, O., Ben-Zur, T., Offen, D., Michaelson, D.M., 2016. Reversal of ApoE4-Driven Brain Pathology by Vascular Endothelial Growth Factor Treatment. Journal of Alzheimer’s disease: JAD 53, 1443–1458.

63. Shi, Y., Holtzman, D.M., 2018. Interplay between innate immunity and Alzheimer disease: APOE and TREM2 in the spotlight. Nature reviews. Immunology 18, 759–772.

64. Shi, Y., Yamada, K., Liddelow, S.A., Smith, S.T., Zhao, L., Luo, W., Tsai, R.M., Spina, S., Grinberg, L.T., Rojas, J.C., Gallardo, G., Wang, K., Roh, J., Robinson, G., Finn, M.B., Jiang, H., Sullivan, P.M., Baufeld, C., Wood, M.W., Sutphen, C., McCue, L., Xiong, C., Del-Aguila, J.L., Morris, J.C., Cruchaga, C., Fagan, A.M., Miller, B.L., Boxer, A.L., Seeley, W.W., Butovsky, O., Barres, B.A., Paul, S.M., Holtzman, D.M., 2017. ApoE4 markedly exacerbates tau-mediated neurodegeneration in a mouse model of tauopathy. Nature 549, 523–527.

65. Sun, B.L., Wang, L.H., Yang, T., Sun, J.Y., Mao, L.L., Yang, M.F., Yuan, H., Colvin, R.A., Yang, X.Y., 2018. Lymphatic drainage system of the brain: A novel target for intervention of neurological diseases. Progress in neurobiology 163–164, 118–143.

66. Sweeney, M.D., Montagne, A., Sagare, A.P., Nation, D.A., Schneider, L.S., Chui, H.C., Harrington, M.G., Pa, J., Law, M., Wang, D.J.J., Jacobs, R.E., Doubal, F.N., Ramirez, J., Black, S.E., Nedergaard, M., Benveniste, H., Dichgans, M., Iadecola, C., Love, S., Bath, P.M., Markus, H.S., Salman, R.A., Allan, S.M., Quinn, T.J., Kalaria, R.N., Werring, D.J., Carare, R.O., Touyz, R.M., Williams, S.C.R., Moskowitz, M.A., Katusic, Z.S., Lutz, S.E., Lazarov, O., Minshall, R.D., Rehman, J., Davis, T.P., Wellington, C.L., Gonzalez, H.M., Yuan, C., Lockhart, S.N., Hughes, T.M., Chen, C.L.H., Sachdev, P., O’Brien, J.T., Skoog, I., Pantoni, L., Gustafson, D.R., Biessels, G.J., Wallin, A., Smith, E.E., Mok, V., Wong, A., Passmore, P., Barkof, F., Muller, M., Breteler, M.M.B., Roman, G.C., Hamel, E., Seshadri, S., Gottesman, R.F., van Buchem, M.A., Arvanitakis, Z., Schneider, J.A., Drewes, L.R., Hachinski, V., Finch, C.E., Toga, A.W., Wardlaw, J.M., Zlokovic, B.V., 2019. Vascular dysfunction-The disregarded partner of Alzheimer’s disease. Alzheimer’s & dementia: the journal of the Alzheimer’s Association 15, 158–167.

67. Thul, P.J., Akesson, L., Wiking, M., Mahdessian, D., Geladaki, A., Ait Blal, H., Alm, T., Asplund, A., Bjork, L., Breckels, L.M., Backstrom, A., Danielsson, F., Fagerberg, L., Fall, J., Gatto, L., Gnann, C., Hober, S., Hjelmare, M., Johansson, F., Lee, S., Lindskog, C., Mulder, J., Mulvey, C.M., Nilsson, P., Oksvold, P., Rockberg, J., Schutten, R., Schwenk, J.M., Sivertsson, A., Sjostedt, E., Skogs, M., Stadler, C., Sullivan, D.P., Tegel, H., Winsnes, C., Zhang, C., Zwahlen, M., Mardinoglu, A., Ponten, F., von Feilitzen, K., Lilley, K.S., Uhlen, M., Lundberg, E., 2017. A subcellular map of the human proteome. Science (New York, N.Y.) 356.

68. van Steensel, M.A., Damstra, R.J., Heitink, M.V., Bladergroen, R.S., Veraart, J., Steijlen, P.M., van Geel, M., 2009. Novel missense mutations in the FOXC2 gene alter transcriptional activity. Human mutation 30, E1002–1009.

69. Verheijen, J., Sleegers, K., 2018. Understanding Alzheimer Disease at the Interface between Genetics and Transcriptomics. Trends in genetics: TIG 34, 434–447.

70. Winchester, L.M., Powell, J., Lovestone, S., Nevado-Holgado, A.J., 2018. Red blood cell indices and anaemia as causative factors for cognitive function deficits and for Alzheimer’s disease. Genome medicine 10, 51.

71. Xian, X., Pohlkamp, T., Durakoglugil, M.S., Wong, C.H., Beck, J.K., Lane-Donovan, C., Plattner, F., Herz, J., 2018. Reversal of ApoE4-induced recycling block as a novel prevention approach for Alzheimer’s disease. eLife 7.

